# Frequency modulation increases the specificity of time-resolved connectivity: A resting-state fMRI study

**DOI:** 10.1101/2023.06.20.545786

**Authors:** Ashkan Faghiri, Kun Yang, Koko Ishizuka, Akira Sawa, Tülay Adali, Vince Calhoun

**Affiliations:** Tri-Institutional Center for Translational Research in Neuroimaging and Data Science (TReNDS), Georgia State University, Georgia Institute of Technology, and Emory University, Atlanta, GA, USA; Department of Psychiatry, Johns Hopkins University School of Medicine, Baltimore, MD, USA; Johns Hopkins University School of Medicine and Bloomberg School of Public Health, Johns Hopkins Medical Institutions, Baltimore, MD, USA; University of Maryland, Baltimore County, Baltimore, MD, USA; School of Electrical and Computer Engineering, Georgia Institute of Technology, Atlanta, GA 30302, USA

## Abstract

The human brain is a highly dynamic system, and the methods we use to analyze the data gathered from this organ should account for this dynamism. One such family of methods that has attracted a lot of attention in the past decades is based on networks. The most well-known method for estimating the connection among these networks uses the sliding window Pearson correlation (SWPC) estimator. Although quite a useful tool, there are some important limitations. One such limitation is that SWPC applies a high pass filter to the activity time series. If we select a small window size (which is desirable to estimate rapid changes in functional connectivity), we will filter out important low-frequency activity information. In this work, we propose an approach based on single sideband modulation (SSB) in communication theory, which aims to solve this issue, allowing us to select smaller window sizes and capture rapid changes in the time-resolved functional connectivity. We use both simulation and real data to demonstrate the superior performance of the proposed method, SSB+SWPC, compared to classical SWPC. In addition, we compare the temporal recurring functional connectivity patterns between individuals with the first episode of psychosis (FEP) and typical controls (TC) and show that FEP stays more in FNC states that show weaker connectivity across the whole brain. A result exclusive to SSB+SWPC is that TC stays more in a state with negative connectivity between sub-cortical and cortical regions. All in all, based on both simulated data and real data, we argue that the proposed method, SSB+SWPC, is more sensitive for capturing temporal variation in functional connectivity.

## 1. Introduction

The human brain is a highly dynamic and complex system, and our measurements of this organ can capture many aspects of this system. In recent years, network science tools have been used extensively for this purpose with much success in capturing both the complexity (Bassett & Gazzaniga, 2011) and the dynamism (Deco et al., 2011) of the human brain. Typically, to use network science tools, one first has to construct the network itself from the recorded brain data. Depending on both the data and the network construction step, we can have many different types of networks. Using functional data like functional magnetic resonance imaging (fMRI), we can capture blood-oxygenation-level-dependent (BOLD) signals and construct functional time-resolved networks. The step of going from the data to the network is quite challenging as we often do not have direct measurements of the network, and there are many open questions about network reconstruction for functional brain networks (Erhardt et al., 2011; Korhonen et al., 2021). A network consists of a set of nodes and the edges that connect these nodes together. To estimate the nodes from the data we can use methods that are based on prior works and atlases (Biswal et al., 1995; Fox et al., 2006) or use data-driven approaches like independent component analysis (ICA) (Calhoun et al., 2009; Faghiri et al., 2018). Some methods, such as constrained ICA (Du et al., 2017; Lin et al., 2010), blend these two categories. The other element of networks are the edges between different nodes which in the neuroimaging field are often referred to as connectivity (e.g., functional connectivity in the case of functional networks).

It is common to use the term functional network connectivity (FNC) when the nodes themselves, are a type of network. For example, if we use ICA to estimate the nodes, these nodes (often called intrinsic connectivity networks or components) are essentially networks themselves (Erhardt et al., 2011). In this work, we use the term FNC exclusively, but the proposed method can also be used to estimate FC. If our aim is to capture the dynamics of the human brain, we should estimate FNC in a manner that is temporally resolved (trFNC). To estimate trFNC from nodes’ time series, many methods have been proposed. The most well-known method for estimating trFNC is the sliding window Pearson correlation (SWPC) which essentially pairs a sliding window with a sample Pearson correlation (Allen et al., 2014). Another method replaces the sliding window part of SWPC with a filter bank to capture trFNC across all its spectrum (Faghiri, Iraji, et al., 2022). Some methods aim to estimate trFNC in a more instantaneous fashion, like instantaneous phase synchrony (Honari et al., 2021; Pedersen et al., 2017), the multiplication of temporal derivative (Shine et al., 2015), or instantaneous shared trajectory (Faghiri et al., 2020). Some methods do not estimate trFNC directly but aim to model it at different levels, like methods based on hidden Markov modeling (Vidaurre et al., 2017). Interested readers can check the many papers that have reviewed the many existing methods for time-resolved connectivity estimation (Iraji et al., 2021; Lurie et al., 2020).

As mentioned above, SWPC is a widely-used method likely because of both its simplicity and the fact that the Pearson correlation itself has been studied for many years (Rodgers & Nicewander, 1988). Looking at SWPC as a system with its sub-system, we can see that it consists of a high-pass filter which is applied to the inputs (nodes time series) in addition to a coupling function and low-pass filter (Faghiri, Iraji, et al., 2022). The window size and window shape selection in SWPC is equivalent to designing these two filters. By selecting smaller window sizes, the high-pass filter cutoff frequency is increased, meaning that we will remove more low-frequency information in the sample space (or activity domain). This can be problematic as the low-frequency content of resting fMRI is quite important (Lee et al., 2013). To solve this issue, some works have suggested a lower bound on the window size for SWPC. For example, Leonardi et al. have suggested that assuming that resting state fMRI data has a lower frequency bound of 0.01, we need to select window sizes larger than 100 seconds to ensure that the high-pass filter in SWPC does not remove important low-frequency information (Leonardi & Van De Ville, 2015). This statement is accurate but with an important caveat. The window size in SWPC also affects its low-pass filter which is applied in the connectivity domain. If we chose a large window size, the low-pass filter’s cutoff decreases, meaning we might smooth out important information in the connectivity time series. It is important to note that we do not have a direct measurement of the connectivity time series; instead, we estimate this time series from the activity time series. So, while there are many studies on the frequency profile of activity time series (Biswal et al., 1995; Chen & Glover, 2015; Fransson, 2005; Trapp et al., 2018) it is quite challenging to discuss the connectivity frequency profile. The challenging part is that we cannot measure connectivity directly and we have to rely on estimators which have their own transfer function that would impact the spectral properties of the connectivity (Faghiri, Iraji, et al., 2022). This is a circular problem: we need to examine the frequency profile of the connectivity time series for selecting the window of the SWPC method but to get the frequency profile of connectivity, we need to use SWPC with a specific window which in turn impacts the estimated frequency profile.

In this work, inspired by a modulation scheme in communication theory we propose a method that allow us to select window size for SWPC without worrying about removing important low-frequency information from the rsfMRI data (Faghiri, Adali, et al., 2022). We evaluate the proposed method using both simulated and real data in addition to using it to analyze a rsfMRI data set

## 2. Methods

### 2.1. Sliding window Pearson correlation (SWPC)

SWPC is an extension of the classical Pearson correlation estimator, where we can capture the correlation coefficients of different temporal segments using a sliding window. The SWPC can be formulated as:

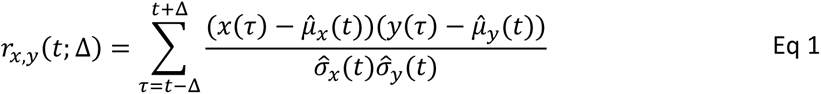

where 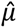_x_(t) and 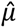_y_(t) are average of x(t) and y(t) for t ∈ [t − Δ, t + Δ] while 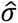_x_(t) and 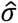_y_(t) are standard deviation of x(t) and y(t) respectively for the range t ∈ [t − Δ, t + Δ]. Note that Δ is the free parameter for Eq 1, which is why it is written after a semicolon.

As discussed in our previous work (Faghiri, Iraji, et al., 2022), the SWPC can be broken into three sub-systems. The first sub-system which both inputs pass through is the part where the moving average is subtracted from the data (i.e., x(τ) − 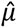_x_(t) in Eq 1 where 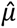_x_(t) is the moving average of x(t)). It is trivial to show that removing the moving average of a time series from itself is equivalent to passing the time series (i.e., x(t) and y(t)) through a high pass filter. The second sub-system is a coupling sub-system which multiplies the outputs of the previous sub-system. The final sub-system is a low-pass filter applied to the coupled time series. Based on this view, the design of the window (its shape and length) is essentially a filter design step for the two filtering sub-systems. As mentioned previously, one of the shortcomings of SWPC arises because of the first high pass of this method (i.e., x(τ) − 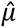_x_(t)). If the window selected for SWPC is small, low-frequency portions of the signals x(t) and y(t) are filtered out which in turn can possibly worsen the estimation of the time-resolved correlation. By selecting large window sizes, we can remedy this issue, but the side effect of such a choice is that we might smooth out important high-frequency time-resolved connectivity information. In this work, we propose a method that allows us to select smaller window sizes for SWPC without removing low-frequency contents of x(t) and y(t) by adding an additional step to the classic SWPC method.

### 2.2. Single-sideband modulation + Sliding window Pearson correlation (SSB+SWPC)

Single sideband modulation (SSB) is a method that can modulate the frequency content of a time series while keeping the signal real. This method can be broken into three different steps. First, the analytic signal is calculated using the Hilbert transform. Second, this complex signal is multiplied with a complex exponential term. And finally, for the third step, the real part of the result of the previous step is calculated. The mathematical formulation of these three steps for x(t) can be simply written as

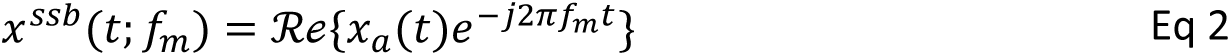

where f_m_ is the modulation frequency and x_a_(t) is the analytic signal calculated using the Hilbert transform x_a_(t) = x(t) + jHx(t)}. Figure 1 demonstrates the steps of SSB in the frequency domain.

**Figure 1.**
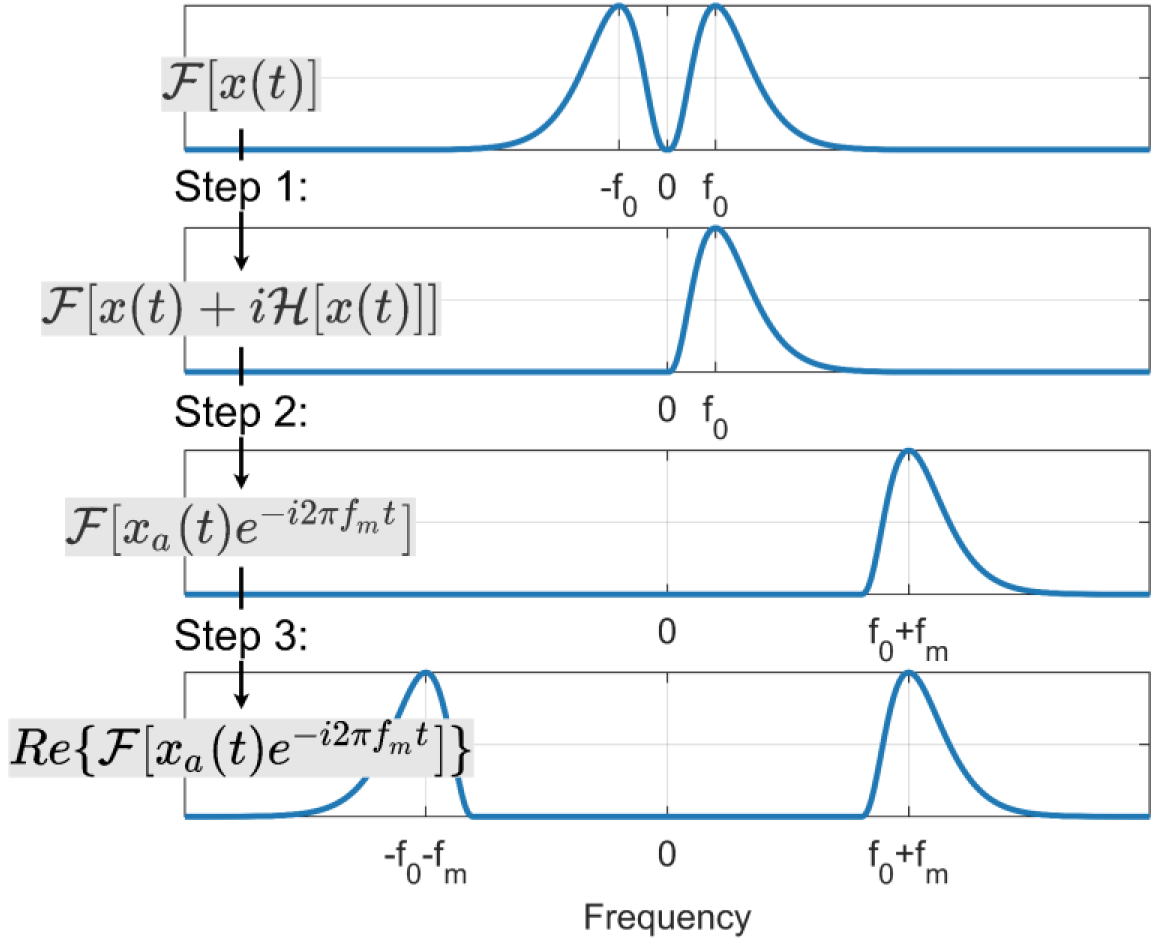
The visual pipeline for the single sideband modulation method. This method has three main steps. In the first step, the analytic signal is calculated using the Hilbert transform. For the second step, the spectrum of the analytic signal is modulated to the right (toward the positive frequencies). In the third step, the negative sideband is added back to the signal by calculating the real part of the modulated signal resulting from the previous step.

By modulating the frequency content of the signal pairs x(t) and y(t) before estimating the SWPC, we can select small window sizes without worrying about filtering out low-frequency information of x(t) and y(t). The formula for this method which we call SSB+SWPC, is:

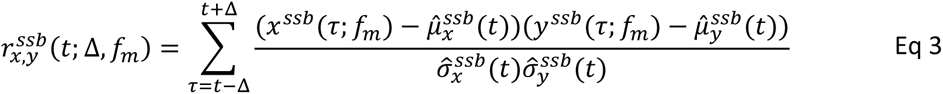

where 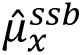 (t) and 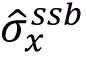 (t) are the moving average and moving standard deviation of the modulated signal x^ssb^(t; f_m_). Note that Δ and f_m_ are the free parameters of the method and are selected by the user. The reason we can use SSB to modulate x(t) and y(t) before estimating the SWPC is because of the spectral property of the coupling sub-function of SWPC. As mentioned before, this sub-function is essentially a multiplication in the time domain (i.e., (x(τ) − 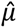_x_(t))(y(τ) − 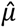_y_(t)) in Eq 1). Multiplication in the time domain is equivalent to circular convolution in the frequency domain (if discrete fourier transform is used for the transformation). And circular convolution can be summarized as flipping and shifting one signal with a specific value and then calculating the area under the curve of the multiplication of the shifted and fixed signal (see Figure 2). We can ignore the flipping step as the spectrums are symmetric. If we modulate x(t) and y(t) using SSB modulation method, the upper and lower sideband of these signals’ Fourier transforms (X^ssb^(f; f_m_) and Y^ssb^(f; f_m_) respectively) are shifted opposite each other to make sure the spectrum remains symmetric (hence why the modulation results are real values).

**Figure 2.**
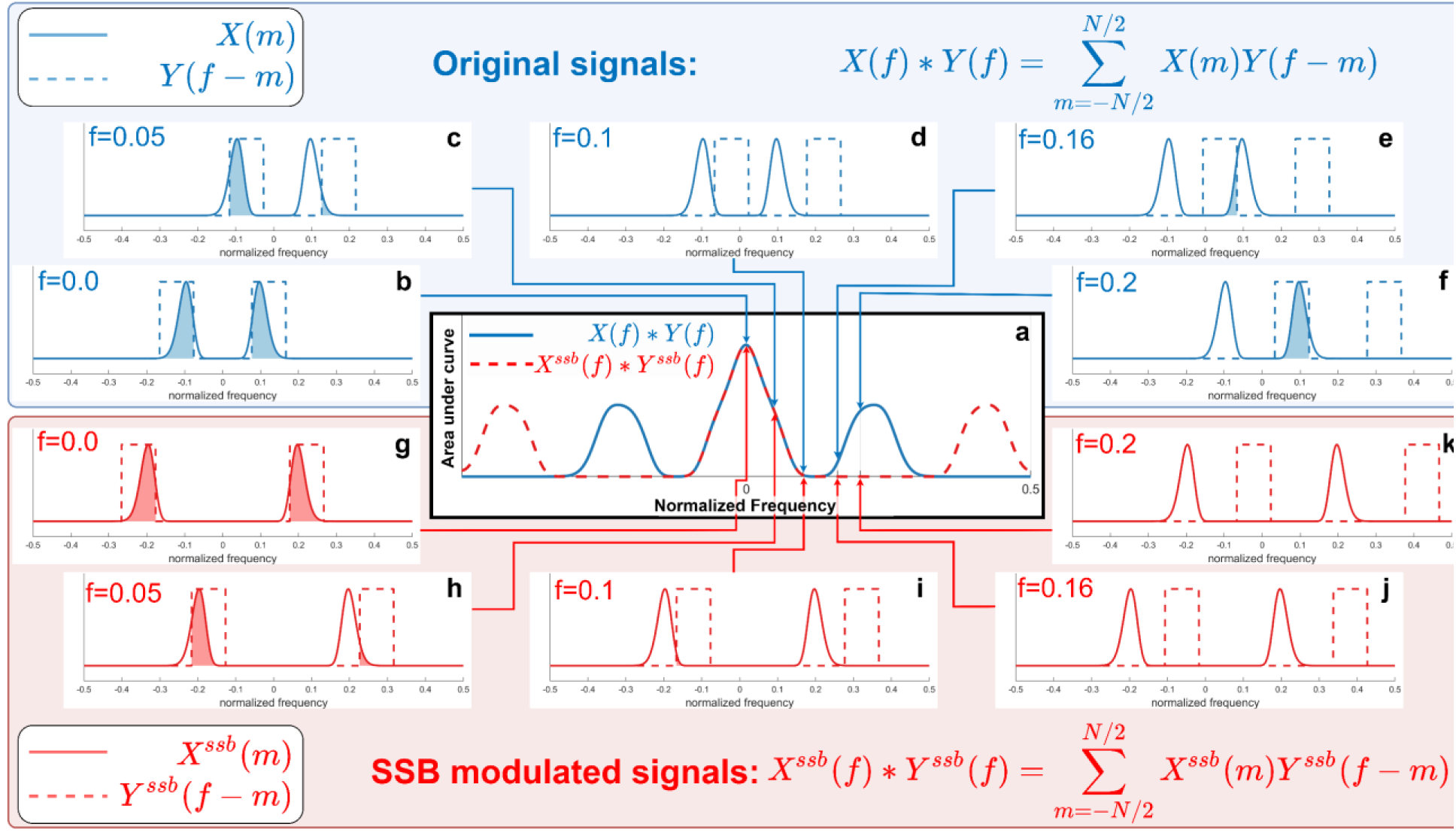
A visual demonstration of why the proposed SSB+SWPC method works. As we saw in the SWPC formula (Eq 1), the coupling part of SWPC is essentially a multiplication in the time domain which translate into a convolution in the temporal domain. Convolution can be viewed as a two step operator Shifting one signal (the dashed lines in this figure) while keeping the other one fixed (the solid lines in this figure), and calculating the area under the curve of the multiplication of these two signals. Therefore as long as the area under the curve for two convolutions is the same, the convolution results are equal for the two operators. Here we see that the area under the curve of the original signal (blue lines) and SSB modulated signals (red lines) are the same as long as the negative sideband of the shifted signal is not overlapping with the positive sideband of the fixed line. i.e., for frequencies below 0.1 Hz, the original convolution results (subfigures b, c, and d) are the same as the SSB modulated signal convolution (subfigures g, h, and i). The two convolutions diverge for shift values higher than 0.1 Hz (subfigures e and f for the original signals and j and k for the SSB modulated signals).

Then the sum of the area under the curve of X^ssb^(f; f_m_) × Y^ssb^(f − f_0_; f_m_) (f_0_ is the frequency shift done as a part of circular convolution) is the same as the area under the curve of X(f) × Y(f − f_0_) for specific shift frequencies f_0_. As you can see in Figure 2 for specific f_0_ values (sub figures b, c, and d for Y(f − f_0_) and g, h, and i for Y^ssb^(f − f_0_; f_m_) in Figure 2). But when the shift value is high enough so that the negative sideband of Y^ssb^(f − f_0_; f_m_) starts overlapping with the positive sideband of X^ssb^(f; f_m_), the two convolution will diverge (see Figure 2a). This value depends on the lowest frequency of the two signals (i.e., X(f) and Y(f)). Meaning that if the two signals have no values at frequencies below a > 0, for shift values lower than 2a, X(f) ⊛ Y(f) and X^ssb^(f) ⊛ Y^ssb^(f) are the same (⊛ is the operator symbol for circular convolution). Contrary to what one might think, this does not mean that X^ssb^(f) ⊛ Y^ssb^(f) have worse estimation for f > 2a, rather, it just means that for these frequencies, X(f) ⊛ Y(f) ≠ X^ssb^(f) ⊛ Y^ssb^(f). As we can see in Figure 2a, the second bump of X(f) ⊛ Y(f) is the result of overlap of positive sideband of X(f) with negative sideband of shifted version of Y(f).

To summarize, SSB+SWPC (the proposed method) adds a modulation step to the classic SWPC method which allows us to select small window sizes for SWPC without worrying about the lower frequency bounds on the activity time series.

### 2.3. Method evaluation using synthesized data

To evaluate the performance of the proposed method, we designed two sets of simulations. The first of these simulations is designed to evaluate if SSB+SWPC improves time-resolved Pearson correlation. The second one is designed to compare a pipeline that uses SSB+SWPC and one that uses classic SWPC in their performance regarding the estimation of states.

For the first set of simulations, we first generated two random time series. Next, we filtered these time series to have a specific bandwidth. For this filtration, we used a Chebyshev Type 2 low pass filter (with the lowest possible order to achieve a maximum of 3 decibels attenuation in the passband and a minimum of 30 decibels attenuation in the stopband). Assume we call these filtered time series x(t) and y(t). Next, we generated connectivity time series to have a specific frequency by using the cosine function C(t) = 0.7cos (2πf_corr_t). For this simulation, it was assumed that variance is constant throughout time and is equal to one. In this case, the covariance matrix is:

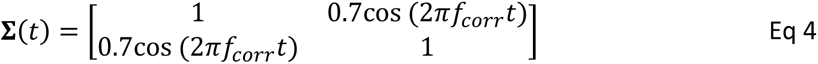

Next, we used Cholesky decomposition to project the two random uncorrelated time series x(t) and y(t), into new time series that are correlated. In other words, we first decomposed the covariance matrix Σ(t) at each time point using Cholesky decomposition **ɛ**(t) = **U**(t)^T^**U**(t) where **U**(t) is an upper triangle matrix. By multiplying **U**(t) with the vector [x(t), y(t)] we will have new time series [x_c_(t), y_c_(t)] that are correlated at each time point according to C(t) = 0.7cos (2πf_corr_t):

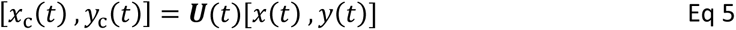

We then used both the proposed method, SSB+SWPC, and the vanilla SWPC approach to estimate time-resolved Pearson correlation (i.e., ^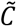^(t)) and evaluated the performance of the two estimators by calculating the Pearson correlation between the estimated correlation (i.e., ^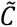^(t)) and the true correlation (i.e., C(t)). This simulation has four free parameters. There are two parameters for the data generation phase, namely the cutoff frequency of the Chebyshev low-pass filter (which determines the true bandwidth of the time series) and f_corr_ (which determines how fast connectivity changes). There are also two free parameters for the analysis: the window sizes for both SSB+SWPC and SWPC methods and the modulation frequency used for SSB.

For the second set of simulations, our goal was to evaluate the performance of SSB+SWPC when it is used in a pipeline to estimate connectivity states of the data assuming we know the true number of the states. This pipeline is essentially the same pipeline we use to estimate the connectivity state of the fMRI data later. We first start by generating six random uncorrelated time series (each with 1000 time points). Then filter them using low pass Chebyshev Type 2, similar to the previous simulation. Next, we generate two six by six matrices to act as the connectivity states. For the first state, the first three time series are highly correlated, while the correlation is zero elsewhere. The last three time series are connected for the second state, while the correlation is zero elsewhere. The correlation amplitude is 0.7 for all connected time series. In other words, the covariance matrix for the two states is:

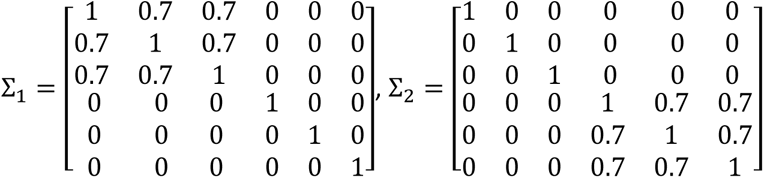

Next, we partition the uncorrelated time series into segments based on the state length. For example, if the state length is 10 samples, we have 10 time points in state 1, followed by 10 time points in state two periodically until we have segmented the whole temporal length of the time series. For each temporal segment, we multiply the two time series by the Cholesky decomposition of the two states’ covariance matrix. Therefore, in that specific segment, we have time series that have correlation matrices depending on the state of the mentioned state. To estimate these correlation matrices, we first use both SSB+SWPC and SWPC methods to estimate connectivity between the 6 generated time series. Next, we concatenate all the estimated connectivity values into a big matrix (for each method separately) with a size equal to T^′^ by 15 (the number of unique connectivity values at each window which is 6 × 5⁄2) where T^′^ = 1000 − (*window size*) + 1. We used k-means clustering to cluster the estimated connectivity matrices into 2 clusters (i.e., we assume that we know the true number of states for this simulation). We use k-means implemented in MATLAB software which uses the k-means++ algorithm for centroid initialization (Arthur & Vassilvitskii, 2006). The clustering was repeated 20 times with the maximum number of iterations set to 500. To evaluate the performance of the two pipelines, we calculated the correlation between the estimated cluster centroid and the associated true state connectivity matrix. There are three free parameters for this simulation. There is the true state length, window size (for calculating SSB+SWPC and SWPC), and modulation frequency (only for SSB+SWPC).

### 2.4. Method evaluation using real data

Using simulated or synthesized data enables us to evaluate the performance of novel methods in a controlled way, but it is also quite beneficial to test the performance of the method using real data. However, this is often quite challenging, especially for data that is the focus of this paper, namely the resting state fMRI data set. To evaluate the performance of any estimator, we need to know the ground truth which we often do not have. In addition, it is unclear how best to compare the results of a novel method with an older method for evaluation, as we do not know what a “good” result looks like. Note that we believe that the goal of a good estimator should be to provide an estimation as close as possible to the ground truth, not to provide estimations that can result in better comparison results between group A and group B using the estimated values. And this is why evaluation using real data is challenging for resting state fMRI connectivity studies. We often do not know what, a good network looks like. Here we come up with a metric that we believe can be used to evaluate trFNC estimation methods at least regarding one specific aspect. First, we argue that trFNC estimation should include time average connectivity (connectivity estimated using the whole temporal range of the time series) information, essentially the information at zero frequency of trFNC. Rather than treating trFNC as a contrast to time-averaged FNC we consider it to be its generalization.

Averaging trFNC gives us its zero frequency (i.e., the time-averaged FNC). Thus, it is desirable for the average of the estimated trFNC to be close to the estimated time averaged connectivity (i.e., connectivity calculated using the whole length of the data). So, the difference between averaged trFNC and time-averaged FNC (using Pearson correlation) can be used as an objective metric to evaluate different methods. So, we first estimate trFNC, using both SSB+SWPC and classic SWPC methods, and then average these matrices across the temporal domain. Then we can calculate the mean square distance between these averaged trFNC and time-averaged FNC (i.e., Pearson correlation using the whole temporal range of the data). The lower this value, the better the performance as this would mean that the estimated trFNC is closer to the true trFNC at zero frequency. Note that this metric only compares the two methods in regard to how well they can capture the zero frequency (i.e., constant) value of the true trFNC and does not provide any information about how well they can capture the temporal variation of trFNC.

### 2.5. Real fMRI data analysis

We used both SSB+SWPC and SWPC to analyze a resting state fMRI dataset. The complete information regarding the dataset used here can be found in previously published works (Faria et al., 2021; Kamath et al., 2019; Kamath et al., 2018). The dataset gathering was approved by the John Hopkins School of Medicine Institutional Review Board (IRB). All participants or their guardians signed a written informed consent. TC and FEP individuals were recruited through the John Hopkins Schizophrenia Center. For an individual to be considered part of the FEP group, they should have had experienced their first episode of psychosis within two years before their enrollment. For this specific study, 183 subjects were included with 94 of those belonging to the TC group. The 89 individuals belonging to the FEP group included SZ (n = 47), schizoaffective disorder (n = 10), schizophreniform disorder (n = 3), bipolar disorder with psychotic features (n = 21), major depressive disorder with psychotic features (n = 5), not otherwise specified psychotic disorder (n = 3).

The fMRI data were prepared for analysis using the statistical parametric mapping (SPM12) toolbox within MATLAB 2019, which included the removal of the first five scans (to ensure signal equilibrium and participants’ adaptation to the scanner). Rigid body motion correction was performed using SPM to correct for subject head motion, followed by slice-timing correction to account for differences in slice acquisition timing. The fMRI data were then transformed into the standard Montreal Neurological Institute (MNI) space using an echo-planar imaging (EPI) template and slightly resampled to isotropic voxels of size 3×3×3 mm³. Gaussian kernel smoothing with a full width at half maximum (FWHM) of 6 mm was applied to the resampled fMRI images. The smoothed datasets were used for the next steps. To decompose the data into a set of spatially maximal independent components, we used an approach based on independent component analysis (ICA) at its core (Calhoun et al., 2009). For this method, first, a brain mask was obtained for each subject by finding all voxels that have higher values (at all time points) than the total average of that subject’s BOLD data. Next, the group brain mask was calculated by finding the voxels that survived in all subjects’ masks. We used the group mask to extract the voxels in the brain for all subjects, normalized the brain data by calculating the z-score, and then applied principal component analysis (PCA) to reduce the temporal dimension of each subject brain data into 120. After concatenating all subjects’ reduced data across their reduced dimension, we applied a second PCA to reduce the dimension of the group data across the concatenated dimension to 100. Finally, we applied an ICA to the PCA results. We used the infomax algorithm for ICA (Bell & Sejnowski, 1995; Lee et al., 1999). To address the issue of the stability of ICA we used the Icasso toolbox and selected the most stable run across the 20 ICA runs (Du et al., 2014; Himberg & Hyvarinen, 2003).

Next, we selected a subset of these 100 components as components of interest. We inspected each component visually and selected components that met the criteria for intrinsic brain networks based on the literature (Allen et al., 2014; Damaraju et al., 2014). we also consider the spectrum of these components’ time series when selecting components (Allen et al., 2011). This resulted in 52 components grouped into seven functional domains. We then use both SSB+SWPC and SWPC to estimate trFNC between all time series pairs. This results in a matrix of size 52×52 for each subject, each window index (i.e., T − 2Δ where T is the total number of time points and the window size is 2Δ + 1). First, we explored the spectral properties of the estimated trFNC using both SSB+SWPC and SWPC methods. Here, we explored the patterns of power spectral density (PSD) of the estimated trFNC. One pattern that has been reported many times for the PSD of brain signals is 1⁄f^β^ (f is frequency, and β is often called power law exponent). This pattern is closely related to a property called scale-freeness which has been reported for many types of signals recorded from the human brain (He, 2011; He, 2014; Lee et al., 2021; Van de Ville et al., 2010). One way we can investigate this property is by first estimating the PSD. We used the function *pspectrum* implemented in MATLAB (The MathWorks Inc. version R2022b) to estimate PSD. We selected 0.5 for leakage (which controls the Kaiser window shape) and 0.01 for frequency resolution bandwidth. After estimating PSD for each subject and connectivity PSD, we fitted a power law function to the PSD using *fit* function of MATLAB with ’power1’ as the model. This resulted in one power law exponent for each subject and connectivity PSD. We compared the power law exponents estimated from SSB+SWPC and SWPC using a two-sample t-test. The *p*-values were corrected for multiple comparisons by controlling for the false discovery rate (FDR) using a linear step-up procedure (Benjamini & Hochberg, 1995).

To investigate the connectivity pattern shared between different subjects at different times, we used a clustering approach. It is important to note that as the connectivity matrices are symmetric, we only have 1326 unique features (i.e., 52 × 51/2) for each matrix. We concatenated the estimated trFNC across all 183 subjects and windows, resulting in a big matrix with 183 × (T − 2Δ) rows and 1326 columns. To summarize these trFNC matrices, we performed clustering using the k-means algorithm (Lloyd, 1982). As the data dimension is high, we used city-block distance (also called Manhattan distance), as suggested by previous works for high dimensional data (Aggarwal et al., 2001). We used the k-means++ algorithm for cluster center initialization (Arthur & Vassilvitskii, 2007). For each cluster number, the clustering was run 20 times with new initial cluster centroids, and the best clustering was chosen based on the run with the lowest within-cluster sum of distances between the points and centroids. To find the best cluster number, we use the elbow criteria on the within-cluster sum of distance. One metric we can calculate based on the clustering results is called dwell time which simply shows how long each subject stays in a given cluster when it goes into that cluster. We have one dwell time value for each subject and each cluster. We compared this metric between TC and FEP groups while controlling for age, gender, and mean framewise displacement. We used robust fitting using fitlm function implemented in MATLAB with ’bisquare’ as the weight function, which has been shown to increase the sensitivity of the neuroimaging analysis (Wager et al., 2005). We corrected for multiple comparisons by correcting for FDR (Benjamini & Hochberg, 1995).

## 3. Results

Two sets of simulations were designed to evaluate the proposed methods in two different scenarios. For the first scenario, we generated two random time series to have a time-resolved correlation that changes as a sinusoid with frequency f_corr_. We then used the proposed method, SSB+SWPC, and classical SWPC to estimate the time-resolved correlation between the time series using different analysis parameters. By comparing the estimated correlation time series with the true time series, we can compare the two methods. We used Pearson correlation as the performance metric here. Figure 3 shows the results for different simulation parameters. Each row shows the results from one specific approach to window selection. The first row shows the results for when the window size is fixed (equal to 5 time points which are quite small), while the second row shows the results for the case where the window size is selected optimally based on the connectivity frequency or f_corr_ (which is unknow for real data). Each column shows the result for a specific f_corr_, while the x-axis of each figure is the modulation frequency (i.e., f_m_ a free parameter of SSB+SWPC). For this specific figure, the two signals’ spectrum had frequency content in the range [0, 0.1] Hz. As we can see here, SSB+SWPC (green line), outperforms SWPC for most frequency modulation values. The performance improvement of using SSB+SWPC is more noticeable if we use a very small window size (5 samples here).

**Figure 3.**
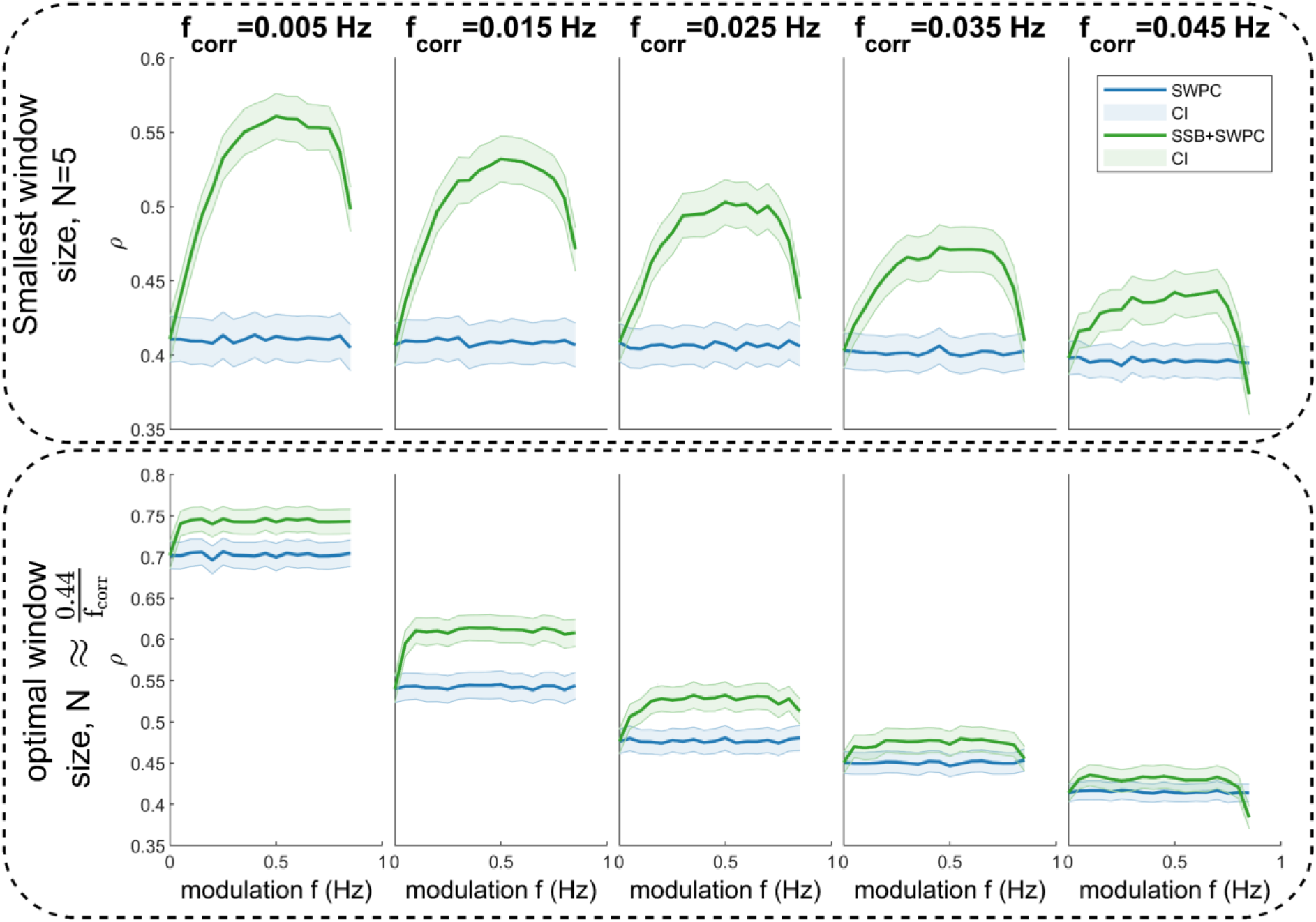
The results for the first set of simulations. Here, we are showing the correlation between the true connectivity values and the estimated values for both SWPC (blue lines) and SSB+SWPC (red lines). The solid lines are the average correlation across all realizations while the shaded area is the 95% confidence interval (CI) of these averages. The first row shows the results of when we chose a very small window size while the second row shows the optimal window size results based on the true connectivity frequency (unknown in real-world settings). Across both scenarios and almost all modulation frequencies, SSB+SWPC outperforms SWPC. The improvement can be as high as p∼0.15 rho which is a substantial improvement.

This is quite logical as using a small window size the high pass filter of SWPC is removing a lot of information from the activity signals in the classical SWPC, while in SSB+SWPC, the low-frequency information is modulated to higher frequencies. If we select the optimal window size (by selecting the window size in such a way that the low pass filter cutoff is equal to the connectivity frequency) we see that the improvement is less pronounced but note that we do not know the frequency of connectivity (as connectivity is the estimated phenomena in the first place), we actually cannot know the optimal window size. Therefore, one can argue that it is quite beneficial to use window small window sizes to be able to estimate connectivities across broader frequency ranges. Another observation we can make based on this figure is that most modulation frequency values result in a better estimation performance, which means that the impact of the choice of this parameter is somewhat lessened. We should just keep in mind to select this parameter in a way that aliasing (modulation of a signal past half of the sampling frequency, i.e., F_s_/2) does not happen. This aliasing is likely why the performance of SSB+SWPC is worse than SWPC for that right-most column in Figure 3 for high modulation frequency values. To summarize, we just make sure that the modulation frequency is less than F_s_⁄2 − b where b is the highest frequency of the activity signals. In resting state fMRI studies, we often pass the activity signals through a bandpass filter. Therefore, this b is determined by the designed bandpass filter.

For the second simulations, we aimed to evaluate the performance of a pipeline that break the data into temporal connectivity states by clustering trFNC using both SSB+SWPC and SWPC method. Here, we examined how well can we estimate the final connectivity states rather than the trFNC value themselves (examined in the previous simulation). This simulation is more relevant as in the actual analysis of the real resting state fMRI data, we break the data into temporal connectivity states and most of the metrics are calculated for those brain states. Each subfigure in Figure 4 shows the final results for one simulation. The rows and columns show the results from different state lengths and window sizes respectively.

**Figure 4.**
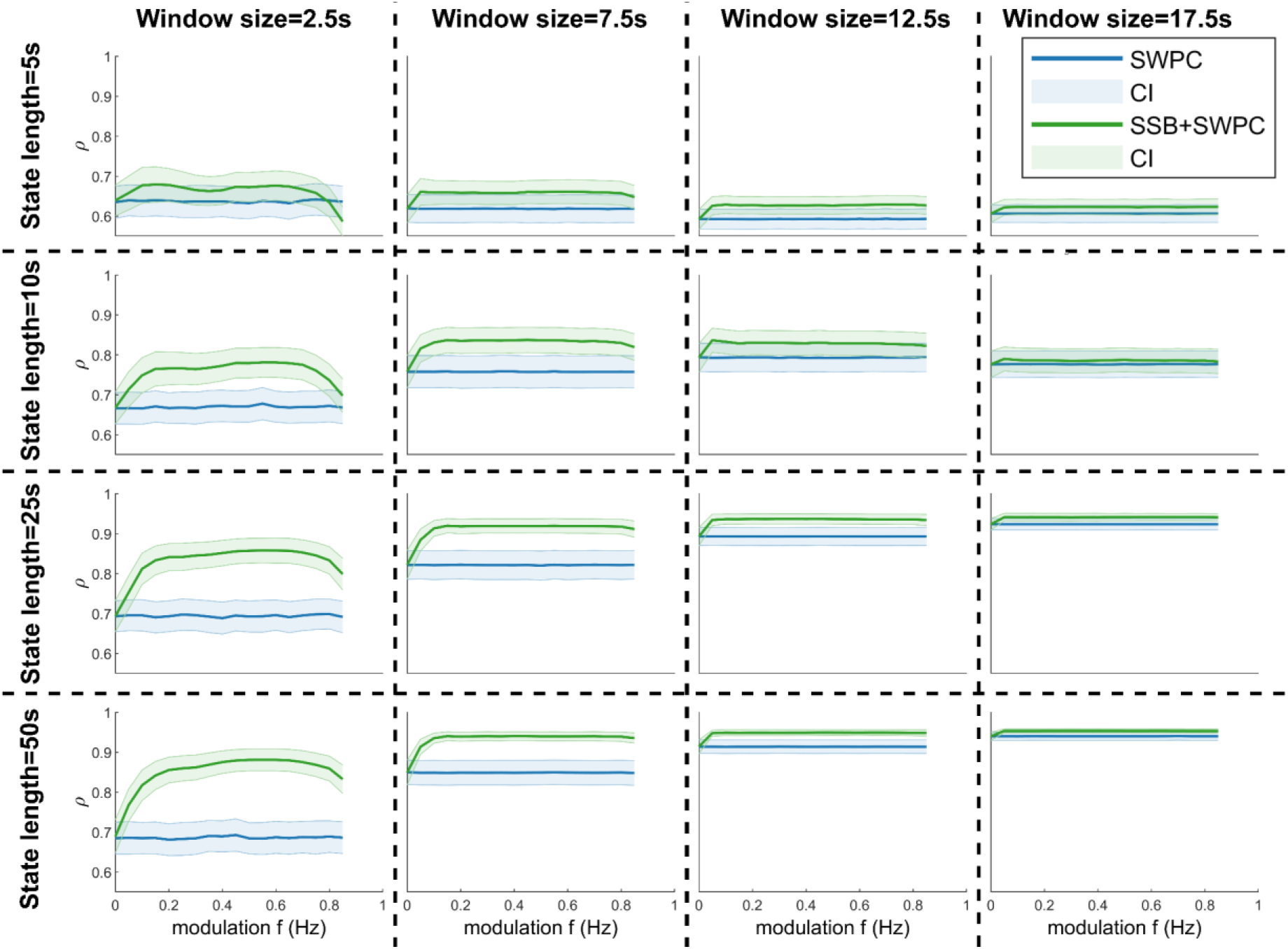
The results of the second set of simulations. Here we are comparing how well we can estimate FNC states using SSB+SPWC (green lines) and SWPC (bluelines) methods. Each row shows the result for one specific state length, while each column shows the result for using one specific window size. We can see that SSB+SWPC outperforms SWPC again across almost all parameters. The performance improvement is more noticeable for smaller window sizes. This makes sense as the larger the window size, the cutoff frequency of the high pass filter of SWPC is lower, meaning we are filtering out less information. Therefore, SSB+SWPC and SWPC results are more similar for larger window sizes.

It can be seen in Figure 4 that using SSB+SWPC improves the performance of the pipeline compared to SWPC across almost all of the different simulation parameters. Similar to the previous simulation, this improvement is more substantial for the smallest window size. For the case of the smallest window size and state length, SSB+SWPC performs worse than SWPC for larger modulation values, but this is likely caused by the aliasing issue we discussed above and can be avoided by choosing an appropriate modulation frequency value. By comparing the results of the first simulation (Figure 3) with this simulation (Figure 4) we can see that the performance is higher across the board for both methods in the second simulation. The reason is that in the first simulation, we are estimating the trFNC values themselves, while for the second simulation, the trFNC values are clustered into a limited number of clusters (two here). Therefore, we are averaging a lot of data to get the states which would mean that some of the noise and error of trFNC estimation are averaged out. Both simulation results showed that using SSB+SWPC we can achieve better performance compared to using classic SWPC in analysis that are based on estimating trFNC.

### 3.1. Real fMRI data analysis

After standard preprocessing, we decomposed the dataset into 100 maximally spatial independent components using the GIFT toolbox and visually selected 52 components based on their spatial maps and spectral properties. These 52 components were then grouped into seven functional domains: Visual (Vis), Somatomotor (SM), Temporal (Temp), Cerebellum (Cb), Default mode (DM), Cognitive control (CC), and Sub-cortical (SC) networks. Figure 5 shows the spatial maps of these 52 components and their grouping. We then used a window size of 7 samples (equal to 14 seconds as TR is equal to 2 seconds) to estimate trFNC using both SSB+SWPC and SWPC methods. The reason we used this short window size was to show the benefit of using SSB+SWPC more, as using a small window size, important low-frequency information will be filtered out in SWPC but not in SSB+SWPC. To get the optimal modulation value for SSB+SWPC, we first calculate the cutoff frequency of the high pass filter of SWPC. As we are using a rectangular window here, the approximate -3dB cutoff frequency of this high-pass filter can be calculated using this formula:

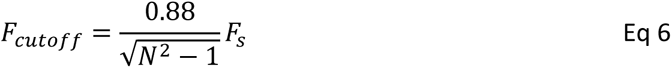

F_s_ is the sampling frequency (i.e., 0.5 Hz) while *N* is the window size (i.e., 7). Using this formula, we get 0.0635 Hz as the cutoff frequency for the high pass filter of SWPC. As the lowest frequency of the activity signals is 0.01 Hz (the lower bound of the bandpass filter applied to the data), we have to modulate the activity signals by 0.625 Hz (i.e., 0.0635 – 0.01) to make sure the high-pass filter of SWPC does not filter important low-frequency information of the activity signals. Next, to summarize the trFNC time series, we applied k-means to cluster the results. To select the cluster number, we used the elbow criteria method. First, we ran the clustering for cluster numbers from 1 to 30 and recorded the sum of square error between cluster centroids and the data in each cluster, and averaged these values across cluster numbers. This resulted in one value for each clustering number. Next, we use elbow criteria to select the best cluster number. This is usually a qualitative step, but here we used a more quantitative method. In short, we fitted two lines to the average within-cluster distance curve. One line fitted to the values belonging to small cluster numbers and one line fitted to the larger cluster numbers. Then the intersection of these two lines can be viewed as the elbow of the curve and be selected as the best cluster number. The intersection of the two lines determines that 4 is the best cluster number based on the elbow criteria.

**Figure 5.**
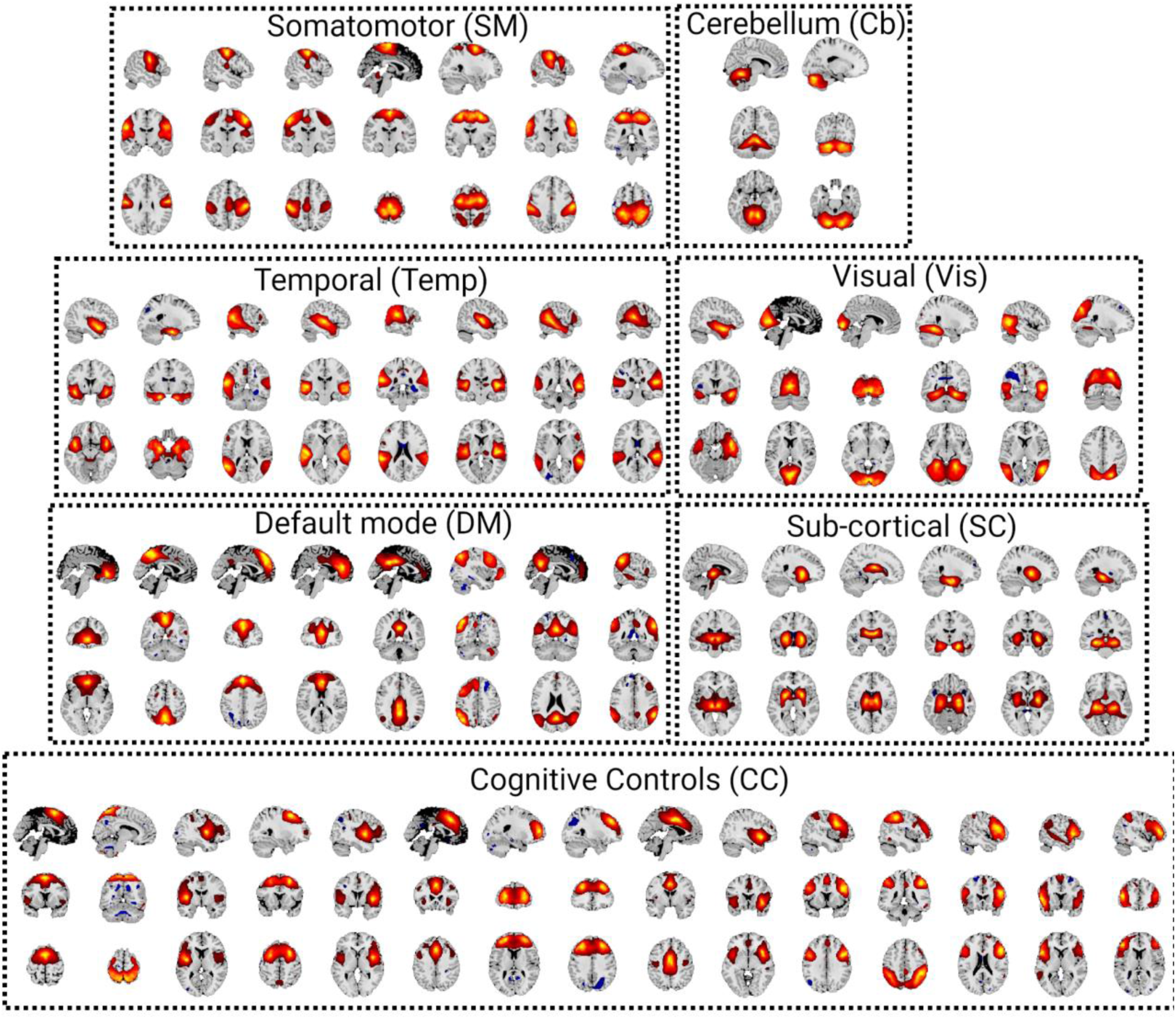
Spatial map of all 52 components grouped into seven functional domains: Somatomotor (SM), Cerebellum (Cb), Temporal (Temp), Visual (Vis), Default mode (DM), Sub-cortical (SC), and Cognitive control (CC) networks

### 3.2. Method evaluation using fMRI data

As mentioned in the method section, we can evaluate a method for estimating trFNC by comparing how well it captures the time-averaged FNC. We can do this by comparing the average trFNC with FNC calculated using the whole temporal range of data using the Pearson correlation estimator (the term static FNC is often used to describe this metric). Figure 6 shows the sFNC for both TC and FEP individuals in addition to the significant (p-value < 0.05) difference between the two groups. We can see here that TC individuals have stronger anti-correlated connections between the sensory functional domains (i.e., visual, sensorimotor, and temporal domains) and cerebellar and subcortical domains. Additionally, we can see that TC has a stronger connection within the temporal functional domain.

**Figure 6.**
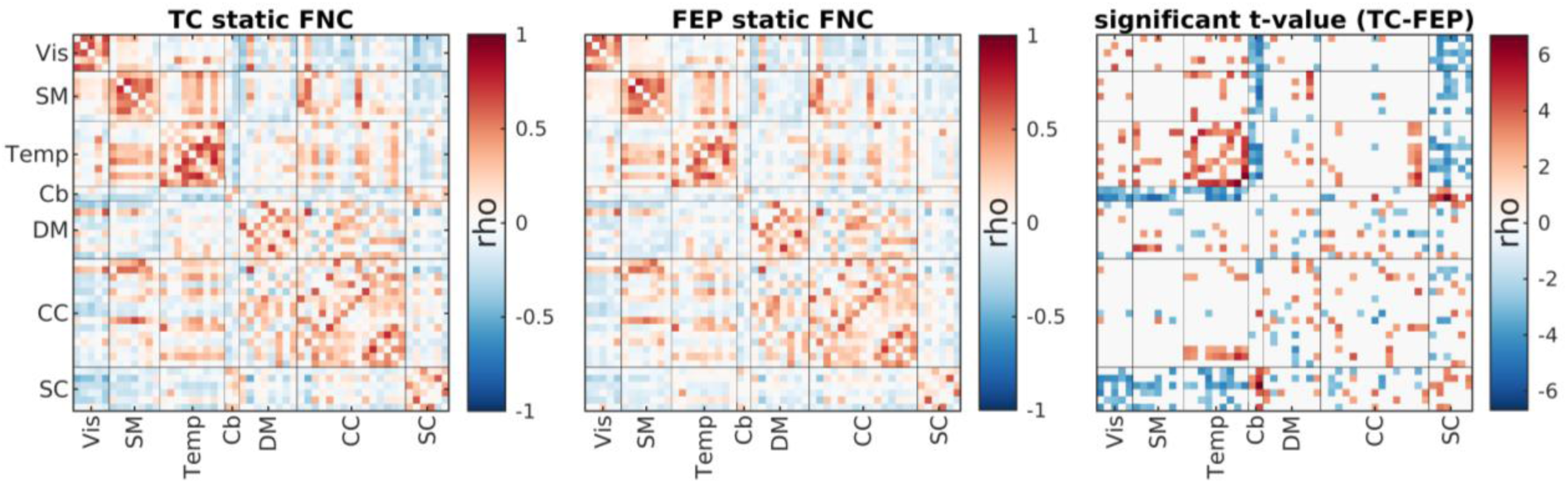
sFNC for both TC (left figure) and FEP (FEP; middle figure). The right figure shows the significant differences between the two groups’ sFNC (corrected for multiple comparisons using FDR). TC individuals have stronger anti-correlated (i.e., more negative) connections between sensory areas (Vis, SM, and Temp) with both Cerebellum and Sub-cortical regions.

The closer the average trFNC and time-averaged FNC are, the better the estimation of trFNC is, at least regarding the content of trFNC at zero frequency. The reason behind this argument is that trFNC is not the opposite of time-averaged FNC (i.e., sFNC) but rather a generalization of it. Time-averaged FNC is essentially an estimate of connectivity at frequency of zero. At the same time, trFNC estimates the connectivity across a broader spectrum, including the zero frequency (at least in the case of SWPC). Therefore it makes sense for us to expect that the average of trFNC estimation using both methods should be close to time-averaged FNC that is estimated directly using Pearson correlation over the whole temporal range of the data. Figure 7 shows the difference between averaged trFNC and time-averaged FNC for both SSB+SWPC and SWPC methods. As we can see, the mean squared error between average trFNC and sFNC is significantly lower for the SSB+SWPC method than the SWPC method (right-most figure), showing that SSB+SWPC outperforms SWPC. This demonstrates that SSB+SWPC can capture the zero frequency content of trFNC better than SWPC. However, this does not tell us how well it can estimate trFNC of frequencies above zero. It is quite challenging to evaluate the performance of any estimator for those frequencies without having access to the true values.

**Figure 7.**
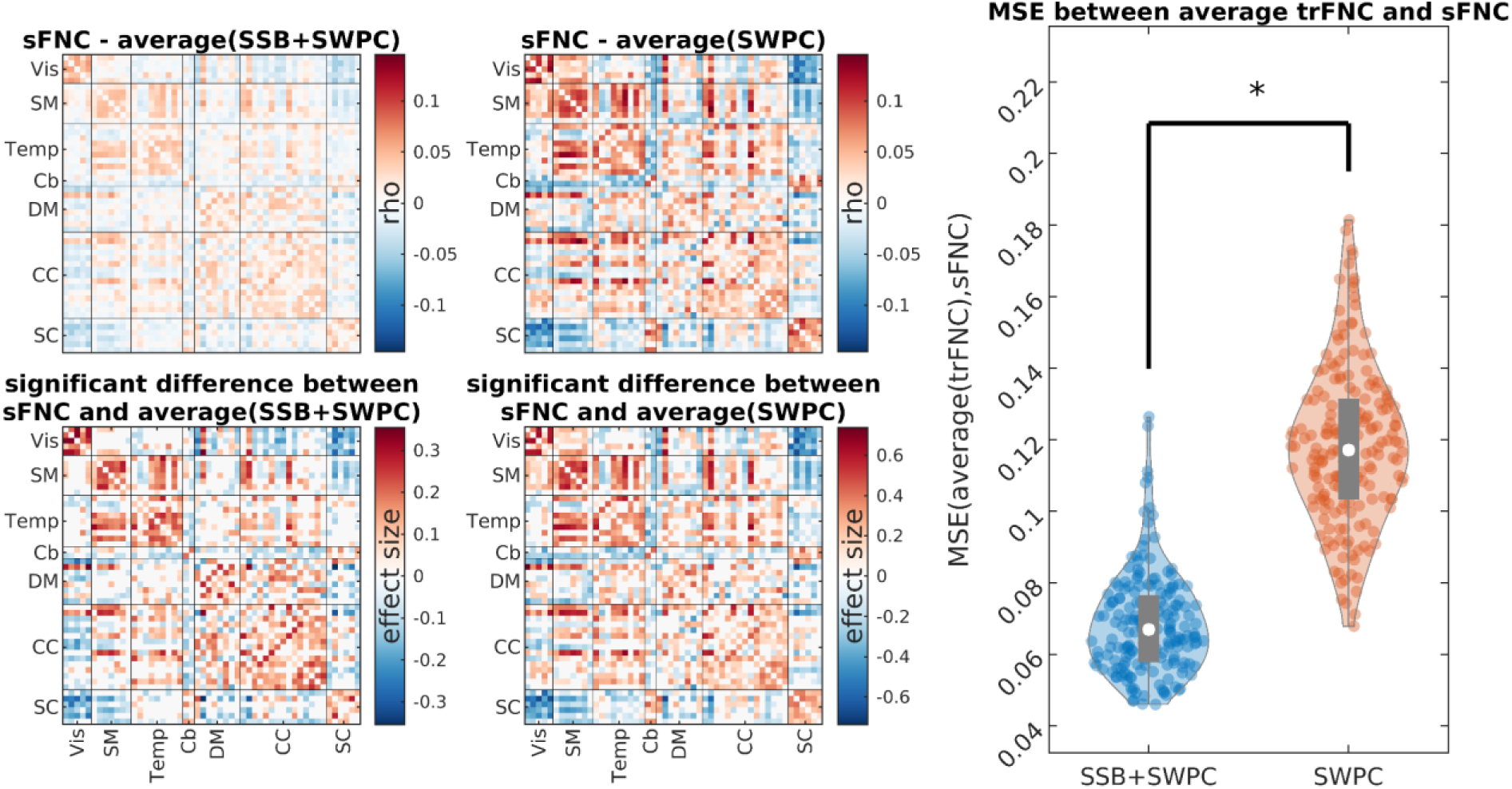
Evaluation of the estimation methods based on how well they can capture time-averaged information. The two figures in the top row left show the difference between averaged trFNC estimation and static (time-average) correlation for both SSB+SWPC and SSB methods, respectively. The bottom row shows the significant (p-value<0.05) differences for the top row. The values in the bottom row are Cohen’s d effect size. The right figures show the MSE between averaged trFNC and time-averaged correlation for the two methods (MSE calculated across all component pairs for each subject). As we can see here, the average of SSB+SWPC is significantly (p-value<0.05) closer to sFNC than the average of SWPC. i.e., SSB+SWPC can capture the zero frequency content of connectivity better than SWPC.

Another metric we can look for a qualitative evaluation of trFNC estimation methods is the spectrum of the estimated trFNC time series. As both methods being compared here, namely, SSB+SWPC and SWPC have a low pass filter as their last sub-system (i.e., the sliding window part), we can compare the average spectrum with the response function of this low pass filter. Ideally, we want a spectrum that is quite similar to the response function of the filter. Figure 8 shows the average spectrum across all subjects and connectivity pairs. As we can see here, SSB+SWPC average spectrum is more similar to the response function of the low pass filter compared to that of SWPC. This is likely because we have selected a very small window size (7 time points) which results in the removal of important low-pass information of the activity signals, which in turn means the signal-to-noise ratio (SNR) is lowered. This does not happen for SSB+SWPC as we have modulated activity signals to higher frequencies so that the low-frequency information is not removed by the high-pass filter portion of SWPC.

**Figure 8.**
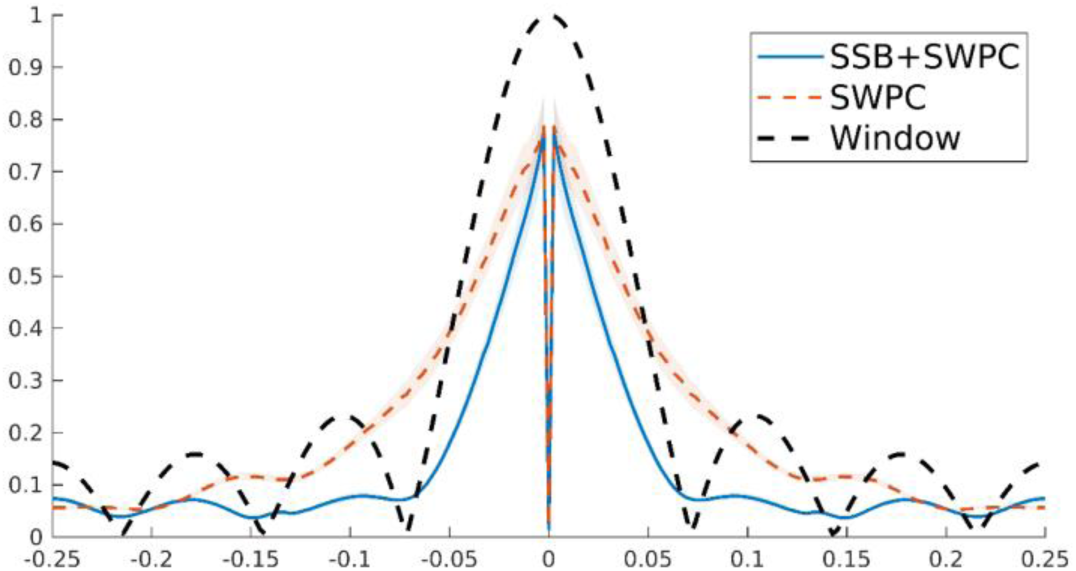
The average spectrum of trFNC estimated using both SSB+SWPC and SWPC methods. As we can see here, the spectrum of the SSB+SWPC estimate follows the transfer function of the sliding window better than estimates of SWPC. Note that as the sliding window is the last sub-system of both methods, the output (i.e., trFNC estimation) should be somewhat similar to the transfer function of the window. This is more important in the sidelobes. We can see here that the SWPC spectrum has high values for at least the first side lobe.

The scale-free property of the trFNC time series is investigated by fitting a power law function (y∼x^β^) to trFNC power spectral density. β is called the power law exponent, and the closer it is to -1, the more complex and scale-free the signal is (Lee et al., 2021). Furthermore, according to The Wiener-Khinchin Theorem, the power spectrum can be understood as the Fourier transform of the autocorrelation function for a wide-sense-stationary random process (Wiener, 1930). If the power spectrum exhibits a steeper slope (indicated by a more negative β value), it indicates a slower and weaker autocorrelation, which would, in turn, mean reduced redundancy. On the other hand, If β is closer to 0, the signal has similar spectral properties to white noise (noise with similar power at all frequencies). Figure 9 shows the average power law exponent for the trFNC time series estimated using both SSB+SWPC and SWPC estimators (the left and middle figures). The right figure shows the difference between the two estimators’ power law exponent values. We can make two observations here; first, we see that intradomain connectivity (especially for Vis, SM, and Temp functional domains) has power exponent values closer to -1 than interdomain trFNCs based on both SSB+SWPC and SWPC results. This might highlight the higher complexity of these specific connectivity time series. The second observation we can make is that trFNC estimated from SSB+SWPC shows a significantly lower power law exponent (i.e., closer to -1) than that of SWPC across many interdomain connections. This might be related to our hypothesis (supported by our simulations) that SSB+SWPC has a higher signal-to-noise ratio (noise here being white noise that has a flat spectrum) as it does not filter out important low-frequency activity information when estimating trFNC.

**Figure 9.**
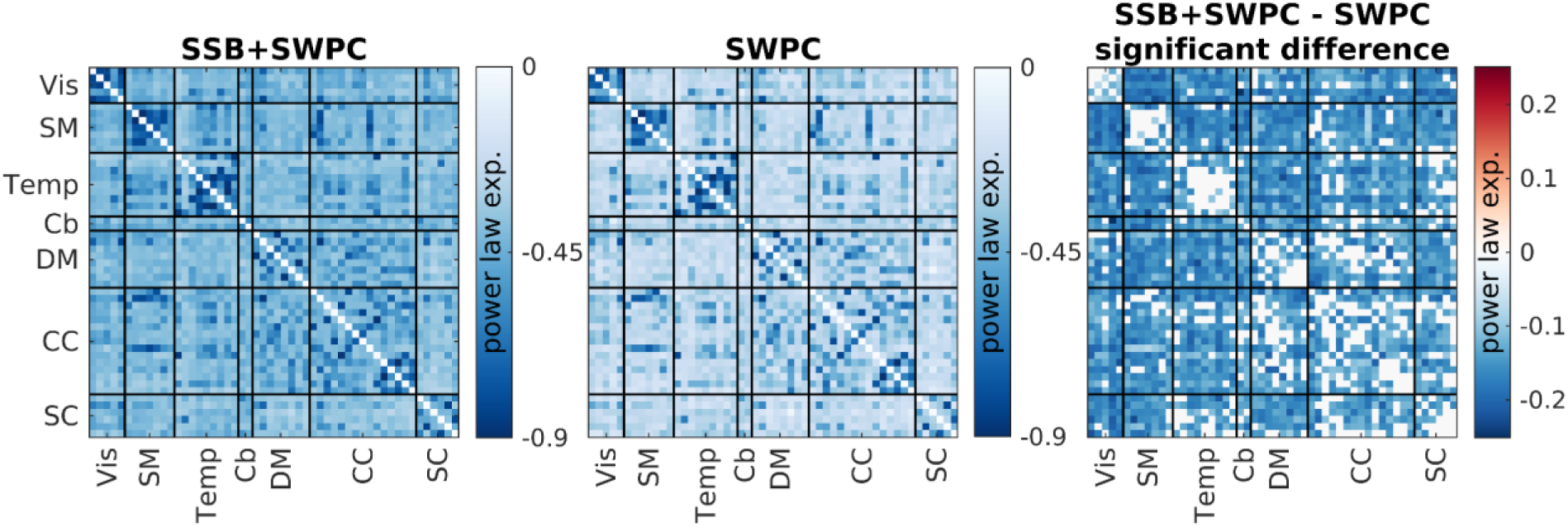
Estimated power law exponent for both SSB+SWPC and SWPC method and their differences. The left figure shows the average power law exponent values (across subjects) for trFNC estimated using SSB+SWPC, while the middle figure shows the same measure for SWPC. The right figure shows the significant difference between the power law components estimated using both methods. A power law exponent closer to -1 might point to a signal with higher complexity, while an exponent closer to zero (i.e., a flatter power spectrum) means that the signal is closer to white noise, at least regarding the power at different frequencies.

### 3.3. Real fMRI data time-resolved results

We can also use the clustering results to compare the two groups (i.e., TC and FEP). One metric we can use for this comparison is the mean dwell time. In short, this value determines how many time points, on average, each subject stays in one cluster once it goes in that cluster. This can be used to measure how much each subject’s data stick to a given FNC state. Figure 10 shows the clustering results for SSB+SWPC while Figure 11 shows the results using SWPC. Based on these two figures, we see that two clusters show significant differences between the two groups’ mean dwell time in SSB+SWPC results.

**Figure 10.**
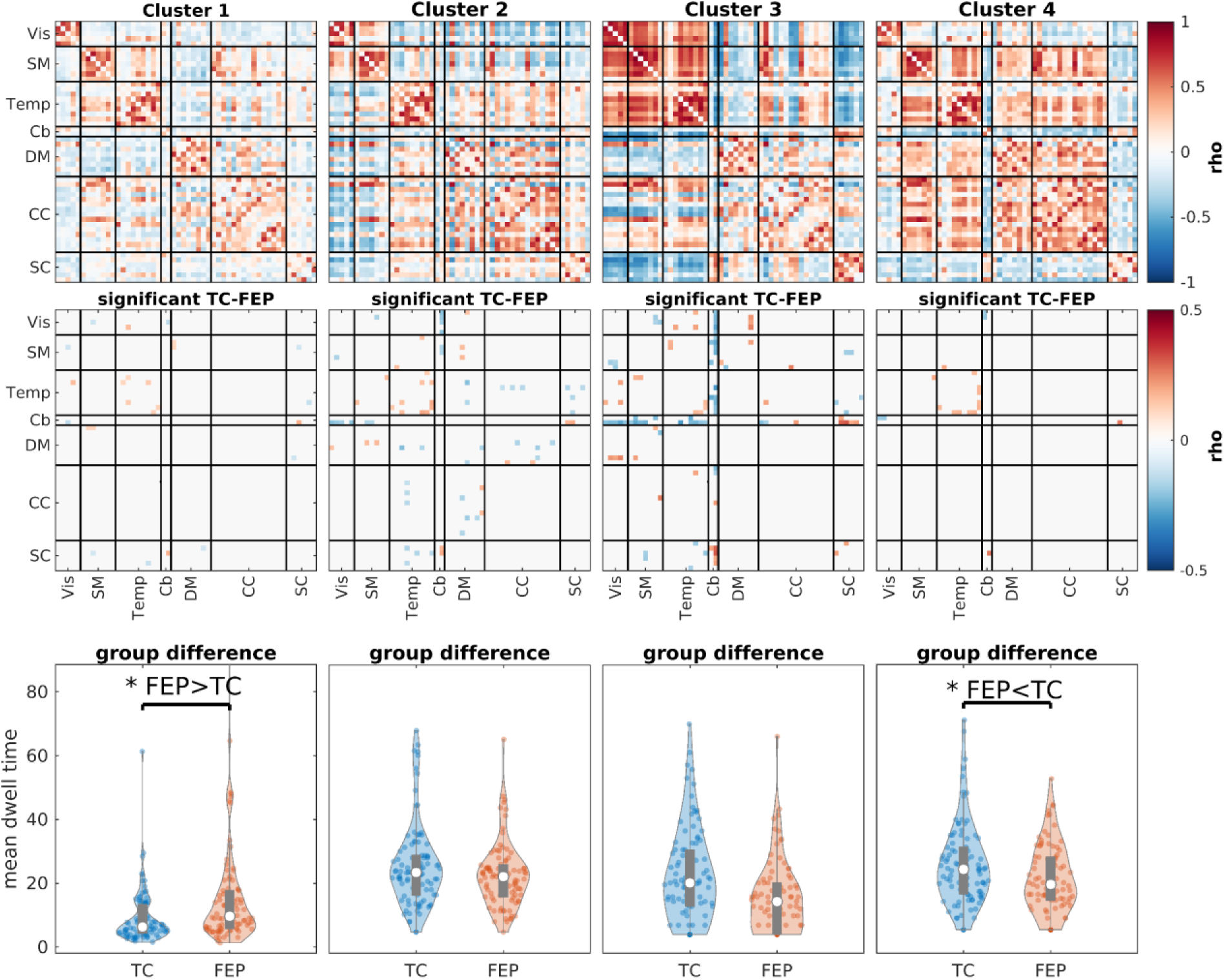
The clustering results for SSB+SWPC. The first row shows the 4 cluster centroids. The second row shows the cells that are significantly different between TC and FEP (subject specific) cluster centroidsThe last row shows the comparison of the mean dwell time for each cluster. Clusters 1 and 4 show significant differences between FEP and TC (p-value<0.05) mean dwell time. FEP individuals generally stay more in state 1, while TC individuals significantly stay more in cluster 4. Cluster 1 shows a more disconnected (i.e., less modular) connectivity pattern (compared to state 4). This is in line with the previous studies that have reported disconnection in individuals with schizophrenia. In addition, TC individuals stay more in state four which is more modular than state 1. State 4 also shows negative inter-domain connectivity between SC (and Cb) and all other functional domains

**Figure 11.**
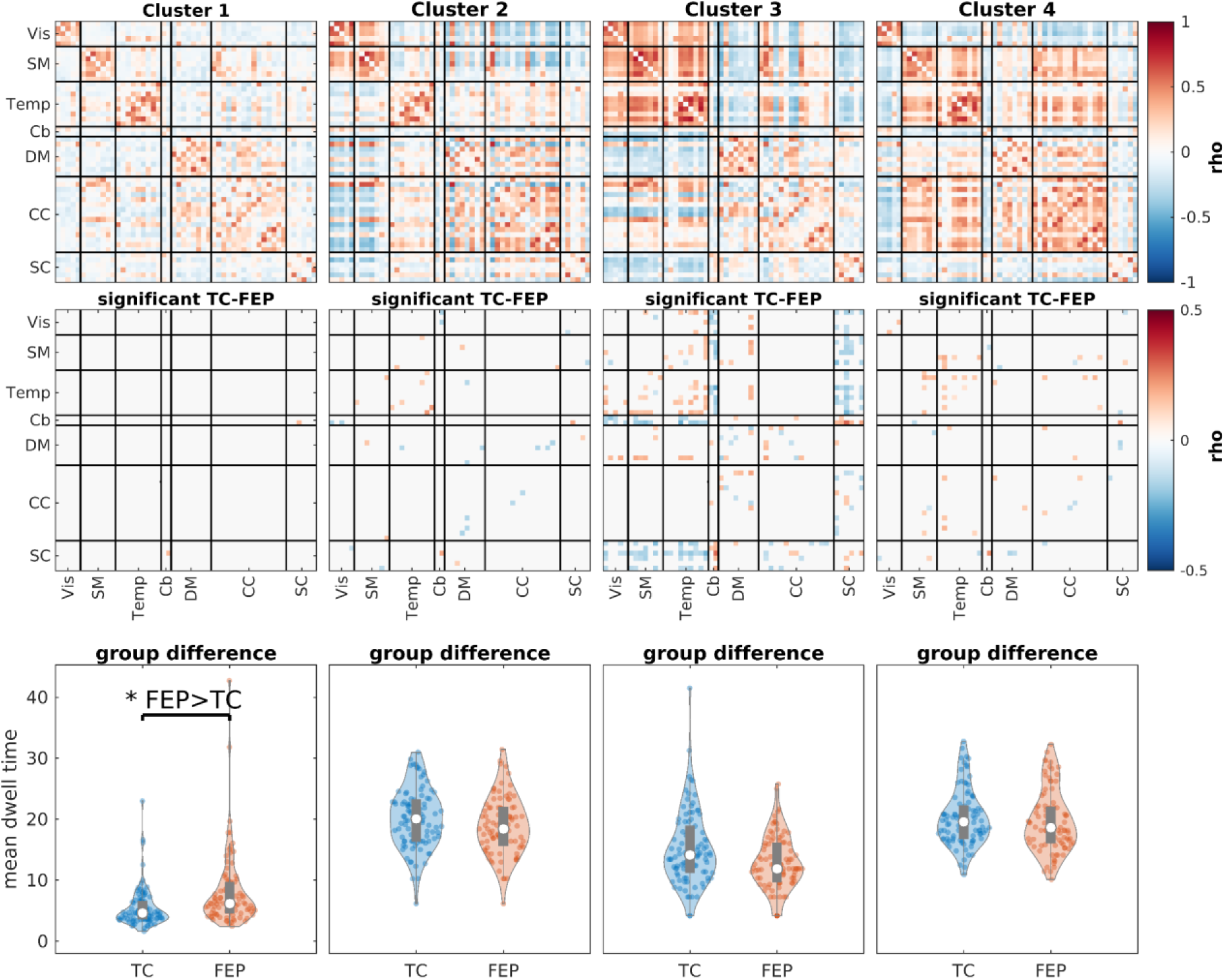
The clustering results for SWPC. The first row shows the 4 cluster centroids. The second row shows the cells that are significantly different between TC and FEP (subject specific) cluster centroidsThe last row shows the comparison of the mean dwell time for each cluster. Similar to SSB+SWPC results (Figure 11), FEP individuals stay more in state 1, which has a weaker connection between different parts of the brain compared to other states. But unlike SSB+SWPC results, cluster 4 dwell time is not significantly different between the two groups.

In contrast, only one cluster shows a significant difference between the two groups for SWPC results. Note that this difference between the two methods’ results should not be considered as us saying that SSB+SWPC is outperforming SWPC. In addition, cluster 4 shows a stronger positive connection between SM/Temp domain and DM/CC domains. And SC components show negative connectivity with all other functional domains other than Cb. These differences are more pronounced in SSB+SWPC results (Figure 10) than in SWPC results (Figure 11).

## 4. Discussion

In this article, we proposed a method to overcome one specific shortcoming of the sliding window Pearson correlation estimator, namely the fact that if a small window size is chosen, some important low-frequency content of the activity signal might be filtered out, resulting in a subpar estimation performance. This is possibly the biggest shortcoming of SWPC, at least for rsfMRI analysis, because of two points: firstly, as we do not know the frequency content of the functional connectivity (most of the research has been on activity spectrum and not the connectivity spectrum), we want to select a small window size so that we can capture the connectivity across a broad range of frequencies. Secondly, much of the important information of rsfMRI activity resides in low frequencies (Biswal et al., 1995; Kalcher et al., 2014; Van Dijk et al., 2010). Although recent studies have increasingly emphasized the presence of rich information in higher frequencies within rsfMRI, challenging our prior understanding (Chen & Glover, 2015; DeRamus et al., 2021; Kajimura et al., 2023), the low frequencies contribute the greatest power it seems. Based on these two points, the method proposed here that allows us to select smaller window sizes for SWPC while improving its estimation can be quite useful. We demonstrated the superior performance of the SSB+SWPC (i.e., the proposed method) compared to the traditional SWPC approach and showcased its application for analyzing a rsfMRI dataset.

For any method aimed at enhancing parameter estimation, it is highly desirable to comprehend why the method outperforms other approaches based on its mathematical formulation. In the method section, we explained why our methods should allow us to use smaller window sizes for SWPC if the data is band limited. In short, the coupling sub-system of SWPC is a multiplication in the time domain, which translates into a circular convolution in the frequency domain.(Faghiri, Iraji, et al., 2022). Therefore, as long as we modulate both input signals of SWPC (i.e., activity signals) with the same frequency value while keeping the spectrum symmetric around zero, the estimated connectivity will not change. Because of this, we can reduce the window size of SWPC without worrying that the high-pass filter of SWPC is filtering out important low-frequency information of the activity signals. This is purely based on the mathematical formulation of SSB+SWPC, and therefore, even without empirical analysis, we can argue that our method should, at least in theory, outperform SWPC. One assumption is that the signal should be band-limited, which is the case for rsfMRI data.

In addition to the theoretical reason why the SSB+SWPC method outperforms SWPC, we also showed the improvement our proposed method offers using two specific simulation scenarios. First, we showed that if our signals are band-limited, and the connectivity is a sinusoid signal with a specific frequency, SSB+SWPC can improve the estimation of time-resolved correlation across most parameters (correlation frequency and modulation frequency). The improvement is pronounced when we use a very small window size, up to ρ∼0.15, which is quite high. But even if we use the optimal window size based on the correlation frequency (not known in real-world settings), SSB+SWPC improves the estimation. The reason behind this is that even if we use the largest window size possible that allows us to capture the connectivity of a specific frequency, this window size might still be smaller than the window size that does not filter out important low-frequency information of the activity signal.

Another way we can view this is that we have two (usually unknown) data parameters that we have to optimize for: the frequency of the activity signal and the frequency of the connectivity signal. But the SWPC method technically has only one parameter, the window size, that we need to select to match the two parameters of the data. The window size should be large enough so that the high-pass filter sub-system of SWPC does not filter out low-frequency information of the activity signal. On the other hand, it should be short enough to capture the connectivity. Depending on the frequency profile of the activity and connectivity signals, these two goals might not be possible with only one parameter. SSB+SWPC, our proposed method, adds an additional analysis parameter in the form of the modulation frequency, allowing us to select the window size without worrying about the activity signal frequency profile. Based on the selected window size, we can modulate the activity signal so that the high-pass filter of SWPC does not filter any important information. In short, the SSB+SWPC has two analysis parameters we can tune to match the data parameter. Note that SWPC technically has another parameter, window shape (e.g., rectangular or Gaussian). But that parameter is minor as it impacts the SWPC filters’ transfer function shape (i.e., the transition between passbands and stopbands) rather than their respective cutoff frequencies most of the time. In addition, we can also have a window shape for SSB+SWPC.

The second simulation showcases how SSB+SWPC performs in a more complete pipeline for analyzing trFNC. Usually, the data we work with has many individual subject data included in it in addition to temporal and spatial information. Many trFNC analysis pipelines aim to first construct networks from these data and then capture a specific number of FNC patterns shared between different subjects at different temporal locations and scales. We often do this with the help of a clustering or dimensional reduction approach. In the second simulation, we assumed that the data occupies a limited number of correlation patterns at different times, often called FNC states in the field. Then we evaluated the performance of a pipeline that uses SSB+SWPC for estimating time-resolved correlation and k-means for the clustering step and compared it with a similar pipeline that uses classic SWPC. We showed that across different parameters of data (correlation state length) and analysis parameters (window size and modulation frequency), SSB+SWPC outperforms SWPC. Similar to the previous simulation, the improvement provided by SSB+SWPC is greater when we use the smallest window size. One observation we made about the result of this simulation is that the performance values are higher than the first simulation for both SSB+SWPC and SWPC methods. The reason for this is that in the first simulation, we are estimating the time-resolved correlation itself, while in the second simulation, we are estimating the handful of shared correlation patterns (i.e., states), meaning we have much more data to estimate a smaller number of values. Therefore, the performance is better across the board. It is important to note that the second simulation is closer to the actual analysis that is often used, as we rarely care about the trFNC values across all time points, nodes, and subjects. We care more about the patterns repeated many times in the data as a whole.

As we mentioned, one assumption of SSB+SWPC is that the signal should be band-limited. So does this mean SSB+SWPC cannot be used for signals with information at every frequency? Not necessary. As we know, if we up-sample a time series that is sampled with the sampling frequency F_s_, by *n*, we will have a new signal with a spectrum in the range [−*n*F_s_⁄2, +*n*F_s_⁄2] but the information is band-limited to the spectral range [−F_s_⁄2, +F_s_⁄2]. Therefore, we can make any signal band-limited by using up-sampling, allowing us to use SSB+SWPC on even white signals. We leave the in-depth exploration of this statement for a future study and only do a simple simulation here. Assume we simulate a pair of time series by drawing 1000 samples from a multivariate normal distribution with mean 0, variance 1, and covariance equal to 0.7 × cos (2π × 0.01 × t). i.e., the correlation changes with time. The sampling frequency is 1Hz. Note that the samples are independent and identically distributed (iid) random variables, so the generated signals have content at all frequencies. Next, we can use SSB+SWPC to estimate the time-resolved connectivity on the original and up-sampled data (up-sampling factor 4). We can evaluate the performance of the estimators by calculating the correlation between true and estimated values. Figure 12 shows the results of a simulation repeated 1000 times for different modulation frequencies, with results of classic SWPC being included at modulation frequency zero. As we can see in Figure 12, SSB+SWPC improves the estimation if it is paired with up-sampling. This improvement is significant across all modulation frequencies, including zero frequency, interestingly. Based on this figure, we see that if we do not use the up-sampling, SSB+SWPC results in worse estimation compared to typical SWPC (results at zero frequency) because of the aliasing that happens at all modulation frequencies. The periodic nature of this result is possibly caused by the fact that the spectrum is cyclic for discrete signals. The dip that happens in the up-sampled results (i.e., green line) is because of aliasing, which was also apparent in Figure 3. To conclude, applying up-sampling on a signal with content at all frequencies enables us to use SSB+SWPC and improve the time-resolved correlation. We leave the more in-depth exploration of this idea to future works, especially because, based on Figure 12, up-sampling can improve classic SWPC results (look at zero modulation frequency).

**Figure 12.**
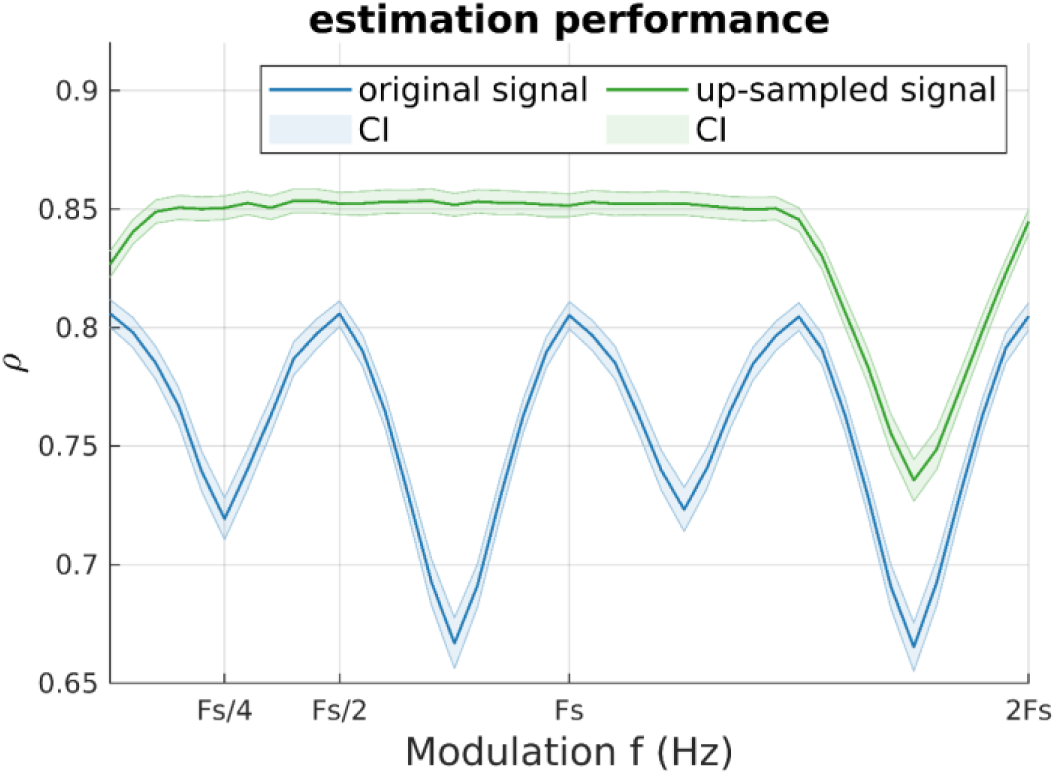
impact of up-sampling on SSB+SWPC estimation performance when the signals have content at all frequencies. The y-axis shows the correlation between the estimand and the true value, meaning higher is better. The blue lines show the correlation for the original signal, while the green line shows the correlation for when up-sampling was used before SSB+SWPC. The shades show the 95% confidence intervals (CI), corrected for multiple comparisons. We can see that using up-sampling allows us to use the estimation improvement provided by SSB+SWPC.

We used a pipeline based on ICA to decompose the preprocessed rsfMRI data from all individuals into 100 spatially maximally independent components. We then selected 52 components and grouped them into 7 functional domains (see Figure 5). trFNC was estimated using both SSB+SWPC and SWPC methods. We first investigated how close the average trFNC was to time-averaged FNC, calculated using the sample Pearson correlation estimator using the whole temporal range of each subject’s data. Figure 6 shows the static FNC For both TC and FEP groups and the significant difference between the groups. We see that TC shows stronger anti-correlation connectivity between Sub-cortical and sensory regions (VIS, SM, and Temp) compared to FEP. Similarly, Cb regions have stronger negative FNC than sensory regions. This aligns with previous research (Damaraju et al., 2014). We also see that TC has stronger connectivity in the temporal domain compared to FEP. Crossley et al. reported dysconnectivity in the temporal lobe (Crossley et al., 2009). We also showed that the average trFNC is closer to sFNC if we use SSB+SWPC significantly in Figure 7, which means that sFNC is captured more accurately in trFNC if we use SSB+SWPC. This is an important point, as the difference between average trFNC and sFNC.

We also explored the spectral properties of the estimated trFNC. Figure 8 depicts the average fast Fourier transform of estimated trFNC using both SSB+SWPC and SWPC methods and the transfer function of the low-pass filter of SWPC. Note that as the last sub-system of SWPC is this low-pass filter, we expect the output (i.e., trFNC) to have a similar spectrum to the transfer function of the low-pass filter. Figure 8 shows that trFNC estimated using SSB+SWPC is more similar to the transfer function than trFNC estimated using classic SWPC. The difference is most apparent in the side lobes of the transfer function, possibly pointing to the fact that SWPC estimates are more noisy. The power spectra of neural activity can provide much meaningful information regarding the human brain (Gao, 2016), but not many studies have explored the spectral properties of the connectivity time series. In this work, we investigated the similarity of the trFNC power spectrum to a power law function (i.e., 1/f^β^). A lot of early studies on different types of signals labeled signals with 1/f power spectrums as noise (Keshner, 1982; Voytek et al., 2015; Zarahn et al., 1997), but more recent papers have argued that signals of interest (e.g., neural activity) can also have 1/f power spectrum (Freeman & Zhai, 2009; He, 2011; He, 2014; Podvalny et al., 2015). Signals with this kind of power spectrum are often called scale-free (He, 2011), which has been reported for many conditions (Ciuciu et al., 2012). E.g., Churchill et al. reported that scale-free extents can be used for measuring cognitive effort (Churchill et al., 2016). We made two observations about the power-law exponent of the results in Figure 9. First, based on both SSB+SWPC and SSB results, we can say that sensory domains (Vis, SM, and Temp) show power exponents that are closer to -1 for intra-domain FNC compared to other inter and intra connections. He et al. reported a power law exponent closer to -1 for activity signals of the visual cortex and -0.6 for the motor cortex (He et al., 2010). Future works must explore the link between the estimated power law exponent of activity and connectivity time series, but based on previous works, we can say that the intra FNC of the sensory functional domains shows higher complexity (Lee et al., 2021). Another observation we can make based on Figure 9 is that the power law exponent is more negative for SSB+SWPC compared to SWPC across many of the connections. The difference is significant for inter functional connectivity power law exponent. As the power spectrum can be interpreted as the Fourier transform of the autocorrelation function (Wiener, 1930), the more negative exponent can be viewed as slower and weaker redundancy.

To compare the reoccurring trFNC patterns between the two FEP and TC groups, we used k-means to cluster the estimated trFNC matrices into 4 clusters. Then we compared the dwell time between the two groups while controlling for nuisance regressors. After controlling for multiple comparisons, we found two clusters that showed significant results for SSB+SWPC (Figure 10) and one for SWPC (Figure 11). Using both methods, we can see that FEP stays significantly more in cluster 1 compared to TC. By visually comparing the centroid for this cluster with other clusters centroids, we can say that cluster 1 shows an FNC pattern that is less connected compared to other centroids. This aligns with the dysconnectivity hypothesis of schizophrenia (Friston et al., 2016). Exclusive to SSB+SWPC result, we can see that TC stays more in cluster 4 on average compared to FEP individuals. First, if we compare cluster 4 to cluster 1, we see a much stronger connection across all regions which again aligns with the dysconnectivity hypothesis of schizophrenia. Additionally, cluster 4 shows strong negative FNC between subcortical regions and cortical regions. Similarly, Chen et al. reported decreased negative connectivity between many cortical regions and the thalamus and reported that (Chen et al., 2019). Another paper reported abnormality in the functional connectivity between the hippocampus and posterior cingulate gyrus, and precuneus for adolescents with early-onset schizophrenia (Wen et al., 2021).

### 4.1. Limitation

The major limitation of the proposed SSB+SWPC method is that it has an additional parameter compared with the classic SWPC method, namely the modulation frequency. The additional parameter gives us more freedom by essentially allowing us to design the low-pass filter of SWPC without worrying about the high-pass filter of the SWPC estimator. But this is a double edge sword as additional parameters increase the complexity and points of failure of methods. For our specific added parameter, the user should make sure to select the modulation frequency in such a way that aliasing does not happen. Another limitation of the SSB+SWPC method is that it assumes that the signals are band-limited. As mentioned previously, we can probably overcome this limitation by up-sampling the time series before using SSB+SWPC, but such a method will introduce at least one additional parameter (i.e., up-sampling factor), which also increases the complexity of the method.

## 5. Conclusion

Our proposed method uses single sideband frequency modulation to mitigate low-frequency information loss even if a small window size is used. We first describe the proposed method, then present several simulation scenarios to evaluate the performance of the proposed method, in comparison with classic sliding window Pearson correlation (SWPC), under specific assumptions. Our simulation results showed that SSB+SWPC is significantly better than SWPC for estimating time-resolved connectivity. We also used the SSB+SWPC method to estimate trFNC from a real resting state fMRI data set to showcase its usage. Even in the real dataset, we showed that SSB+SWPC outperforms SWPC for the estimated time-resolved sample Pearson correlation. Using SSB+SWPC also resulted in significant differences between TC and FEP individuals, consistent with but extending prior work. Overall, the simulation results combined with real fMRI results indicate that the proposed approach (i.e., SSB+SWPC) outperforms SWPC in many scenarios.

## CRediT authorship contribution statement

**Ashkan Faghiri:** Conceptualization, Methodology, Formal analysis, Writing - original draft, Software, Visualization. **Kun Yang:** Writing - review & editing, Resources, Data curation. **Koko Ishizuka:** Writing - review & editing, Resources, Data curation. **Akira Sawa:** Writing - review & editing, Resources, Data curation. **Tülay Adali**: Writing - review & editing. **Vince Calhoun**: Conceptualization, Methodology, Writing - review & editing, Supervision Resources, Funding acquisition.

## Acknowledgment

This work was supported by the National Institutes of Health (NIH) grant (R01MH123610, and the National Science Foundation (NSF) grant #2112455.

## Data and code availability statement

The function for estimating SSB+SWPC in both python and MATLAB language can be accessed through GitHub (https://github.com/afaghiri/SSBSWPC). We have also written a jupyter notebook to act as a demo for this paper and shared it on the GitHub page. Additionally, the function will also be implemented in the GIFT toolbox (https://trendscenter.org/software/). The data were not collected by us and are provided in a deidentified manner. The IRB will not allow sharing as a data reuse agreement was not signed by the subjects during the original acquisition. We can share the derived results as they are not considered human subjects research.

